# Activin receptor ALK4 promotes adipose tissue hyperplasia by suppressing differentiation of adipocyte precursors

**DOI:** 10.1101/2022.10.31.514502

**Authors:** Ee-Soo Lee, Tingqing Guo, Raj Kamal Srivastava, Assim Shabbir, Carlos F. Ibáñez

## Abstract

Adipocyte hyperplasia and hypertrophy are the two main processes contributing to adipose tissue expansion, yet the mechanisms that regulate and balance their involvement in obesity are incompletely understood. Activin B/GDF-3 receptor ALK7 is expressed in mature adipocytes, and promotes adipocyte hypertrophy upon nutrient overload by suppressing adrenergic signaling and lipolysis. In contrast, the role of ALK4, the canonical pan-activin receptor, in adipose tissue is unknown. Here we report that, unlike ALK7, ALK4 is preferentially expressed in adipocyte precursors, where it suppresses differentiation, allowing proliferation and adipose tissue expansion. ALK4 expression in adipose tissue increases upon nutrient overload and positively correlates with fat depot mass and body weight, suggesting a role in adipose tissue hyperplasia during obesity. Mechanistically, ALK4 signaling suppresses expression of CEBPα and PPARγ, two master regulators of adipocyte differentiation. Conversely, ALK4 deletion enhances CEBPα/PPARγ expression and induces premature adipocyte differentiation, which can be rescued by CEBPα knock-down. These results clarify the function of ALK4 in adipose tissue and highlight the contrasting roles of the two activin receptors in the regulation of adipocyte hyperplasia and hypertrophy during obesity.

## Introduction

The contributions of hyperplasia and hypertrophy to adipose tissue expansion during obesity are diet-, sex- and fat depot-dependent. In obese women, hyperplasia was found to be predominant in the subcutaneous fat depot, whereas adipocyte hypertrophy was observed both in the omental (visceral) and subcutaneous compartments (1). In a study on mice fed a high-fat diet, visceral fat grew mostly by hypertrophy and subcutaneous fat by hyperplasia (Joe 2009). Using different strains of mice and mathematical modeling, another study found that hypertrophy is strongly correlated with diet, while hyperplasia was more dependent on genetics (Gavrilova 2009). Yet other studies found that high fat diet promoted proliferation of adipocyte precursors and contributed to adipose tissue hyperplasia in mice, particularly in the visceral fat depot (2, 3). From these and other studies, it transpires that the mechanisms that regulate depot-dependent, diet-mediated adipose tissue growth by hyperplasia vis-a-vis hypertrophy are incompletely understood and remain to be clarified.

Members of the transforming growth factor-β (TGF-β) superfamily of ligands and receptors are potent regulators of cell proliferation and differentiation. Cell culture experiments have shown that TGF-β and activin A negatively regulate pre-adipocyte differentiation (4–7). However, the nature of the ligands and receptors that mediate such effects in vivo and their physiological relevance or requirement, if any, have not yet been elucidated. The activin B and GDF-3 receptor ALK7, a member of the TGF-β superfamily of receptors, is highly expressed in mature adipocytes of both human and mouse (8, 9). Studies in genetically modified mice have shown that ALK7 promotes adipocyte hypertrophy and fat accumulation upon nutrient overload by suppressing lipolysis (10, 11) through downregulation of the expression and signaling of adipocyte β-adrenergic receptors (10), a phenomenon known as catecholamine resistance (12–14). As a consequence, mice lacking ALK7 in adipose tissue are resistant to diet-induced obesity (10, 15, 16). ALK4, a receptor structurally related to ALK7 encoded by the *Acvr1b* gene, is capable of interacting with a much larger ligand repertoire than ALK7, including activin A, to which ALK7 cannot bind (17). The fact that activin A displays effects on fat cells in culture (6, 7) suggests an involvement of this receptor in adipose tissue homeostasis. ALK4 expression is rather ubiquitous, as most cell types, tissues and organs express some level of this receptor. However, the actual cell types that express ALK4 in adipose tissue have not been characterized (9). Moreover, the physiological relevance or function of ALK4 in fat tissue remains to be elucidated.

In this study, we set out to characterize the cellular expression, regulation and function of ALK4 in subcutaneous and visceral adipose tissues through gain- and loss-of-function experiments using ligand activation and ALK4-deficient primary adipocytes, respectively, as well as in vivo studies in mouse strains lacking ALK4 expression from different stages of adipocyte differentiation subjected to diet-induced obesity. Our results show that, in contrast to ALK7, ALK4 promotes adipose tissue hyperplasia by cell-autonomously suppressing differentiation of adipocyte precursors in a diet-dependent manner.

## Results

### ALK4 expression in adipose tissue is induced by caloric intake and dynamically regulated during a dietary cycle

The expression of mRNA encoding ALK4 was investigated in inguinal subcutaneous white adipose tissue (SWAT) and epididymal white adipose tissue (EWAT) in 20 week-old C57BL/6 mice kept on a normal Chow diet (5% of calories from fat) or a high-fat diet (HFD, 60% of calories from fat) during the last 14 weeks. Expression of Alk4 mRNA was significantly increased in both depots under HFD conditions (Fig. 1A). In contrast, the levels of mRNA encoding ALK7 were decreased in the mice fed on HFD (Fig. 1A). As previously reported (Guo 2014), expression of Adrb3 mRNA, encoding the type 3 β-adrenergic receptor, was also downregulated in HFD (Fig. 1A), a phenomenon known as diet-induced catecholamine resistance (12–14). While the level of Inhba mRNA, encoding activin A was unchanged by diet, Inhbb mRNA, encoding activin B, was increased in SWAT after HFD (Fig. 1A), in agreement with previous results in obese patients (9). Finally, expression mRNA encoding GDF-3 was significantly increased under HFD (Fig. 1A), in line with earlier observations (18, 19).

**Figure 1.**
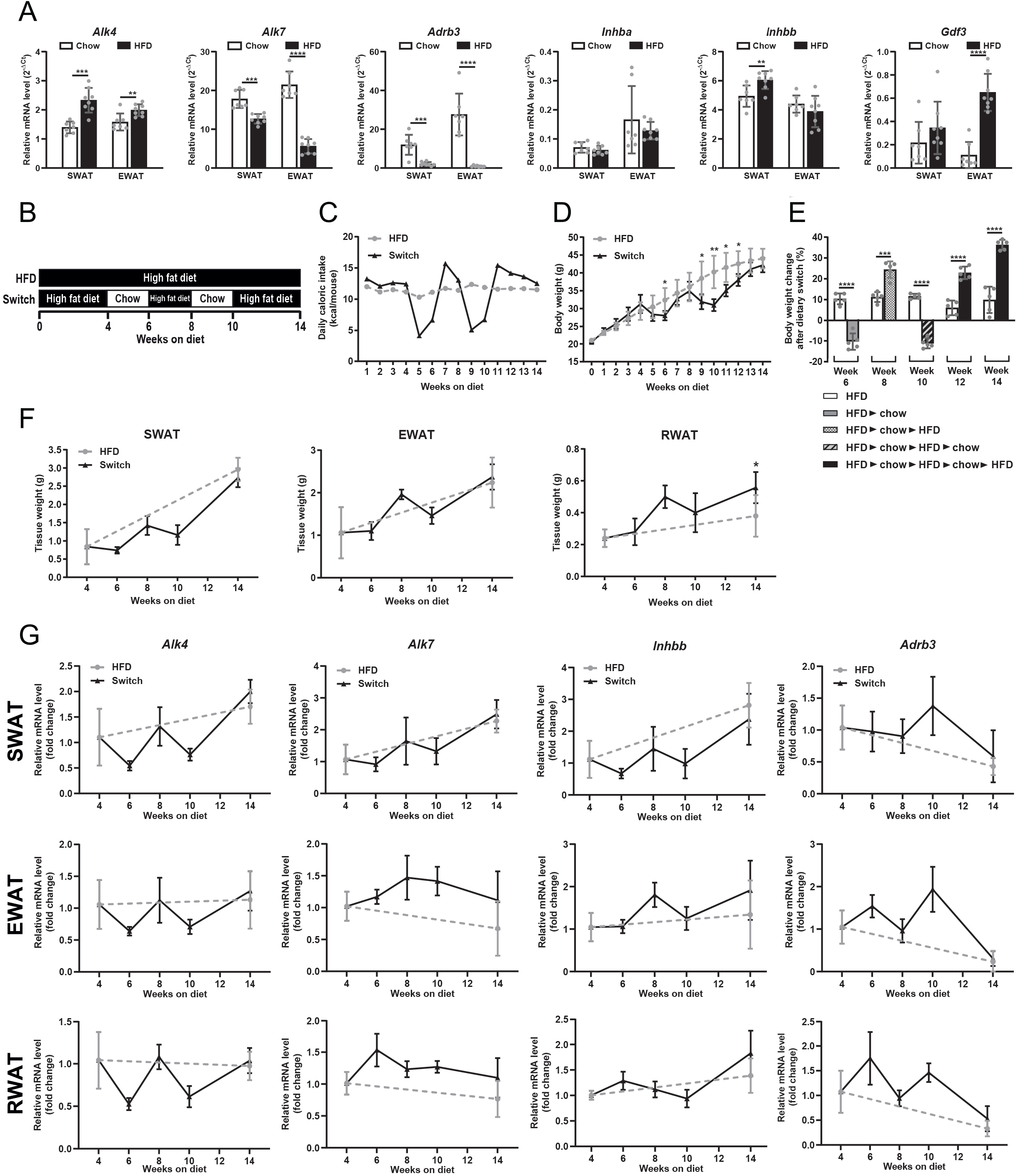
ALK4 expression in adipose tissue is induced by caloric intake and dynamically regulated during a dietary cycle. A) Expression of mRNAs encoding ALK4, ALK7, Adrb3 and ALK4 ligands activin A (Inhba), activin B (Inhbb) and Gdf3 in subcutaneous inguinal white adipose tissue (SWAT), epididymal WAT (EWAT) and retroperitoneal WAT (RWAT) in 20 week old C57Bl/6 mice under standard chow diet (Chow, N=7) and after 14 week high fat diet (HFD, N = 8). mRNA levels were normalized to housekeeping genes Tbp and Ywhaz. Data are presented as mean ± SD. **p<0.01; ***p<0.001; ****p<0.0001 (unpaired t-test). B) Experimental scheme of the dietary cycle. Six week old mice were subjected to a alternating diets for a period of 14 weeks as indicated (Switch). Mice fed a constant HFD served as controls. C) Mean daily caloric intake in mice under constant HFD (grey circles) and alternating diets (Switch, black triangles). D) Weekly body weight change of the diet Switch group (black triangles) in comparison to constant HFD group (grey circles). Data are presented as mean ± SD. *p<0.05; **p < 0.01 (one-way ANOVA followed with Tukey’s Sidak’s multiple comparison test) N=5 mice per group. E) Plot of bi-weekly body weight change in Switch and HFD groups as indicated. Data are presented as mean±SD. ***p < 0.001; ****p < 0.0001 (2-way ANOVA, N=5 mice per group). F) Weights of dissected SWAT, EWAT and RWAT in diet Switch group (black triangles) in comparison to constant HFD group (grey circles). Data are presented as mean ± SD. *p<0.05 (2-way ANOVA, N=5 mice per group). G) Expression of mRNAs encoding ALK4, ALK7, activin A (Inhba) and Adrb3 in SWAT, EWAT and RWAT in diet Switch group (black triangles) in comparison to constant HFD group (grey circles). mRNA levels were normalized to housekeeping genes Tbp and Ywhaz. Data are presented as mean ± SD. N=5 mice per group.

In order to further investigate the contrasting patterns of ALK4 and ALK7 expression in adipose tissue under different diet regimes, we subjected cohorts of C57BL/6 mice to alternate periods of Chow and HFD (Switch) and compared them to a cohort kept under constant HFD (Fig. 1B). The daily caloric intake of mice in the Switch group oscillated as expected according to the diet, while that in the HFD group remained constant (Fig.1C). Notably, the weight gain trajectories of the two groups remained relatively close (Fig. 1D). Thus, while the Switch group lost a few grams during the Chow diet period, it recovered weight at a higher rate when switch back to HFD, as shown by the increased body weight gain following each diet switch (Fig. 1E). At the end of the study, the average weight of the 2 groups was statistically indistinguishable, despite the Switch group spending 4 weeks of the 14 week period under a low-caloric regime. This phenomenon has been previously attributed to metabolic rate adaptation following a period of significant weight loss (Müller 2013, Fothergill 2016, Srivastava 2021). The weights of three different adipose tissue depots, SWAT, EWAT and retroperitoneal white adipose tissue (RWAT) followed a similar oscillatory pattern as predicted by the caloric intake and overall body weight of the animals (Fig. 1F). Remarkably, however, by the end of the experiment, the weight of the RWAT depot ended up to be significantly higher in the Switch group compared to the group maintained under constant HFD (Fig. 1F, right panel). Interestingly, the levels of mRNA encoding ALK4 underwent rather strong oscillations (about 2-fold) in response to the alternative diets in the Switch group in all three fat depots (Fig. G, first column). Inhbb mRNA, encoding activin B, followed a similar pattern, although of somewhat smaller amplitude (FIg. 1G third column). Levels of Adrb3 mRNA also oscillated but in the opposite direction, as expected from the suppressing effects of HFD on Adrb3 expression (Fig. 1G, last column). In contrast, expression of mRNA encoding ALK7 showed a different pattern in mice subjected to alternate diets (Fig. 1G, second column). In SWAT and RWAT, Alk7 mRNA levels showed low amplitude oscillations in opposite directions, while remaining nearly constant in EWAT. We speculate that perhaps 2 weeks may be too short a period to fully reset the transcription levels of this gene upon a caloric intake switch. Nevertheless, together these data reveals contrasting patterns of gene expression in Alk4 and Alk7 in response to different diet regimes.

### Inactivation of ALK4 in adipose tissue from birth attenuates postnatal tissue expansion but has no effect on diet induced obesity in adult mice

In order to begin addressing the function of ALK4 in adipose tissue, we crossed *Acvr1b*^fx^ mice (20)) with *Ap2*^CRE^ mice (21) to generate animals lacking ALK4 expression in adipose tissues from birth. At 4 month of age, *Ap2*^CRE^::*Acvr1b*^fx/fx^ mutant mice kept in Chow diet showed normal body weight (Fig. 2A) and total lean mass (Fig. 2B), but significantly reduced total fat mass (Fig. 2C). Accordingly, leptin serum levels were 8-fold lower in the mutants compared to *Acvr1b*^fx/fx^ controls (Fig. 2D). Macroscopic examination of epididymal adipose tissue revealed profound lipodystrophy in the mutants (Fig. 2E). Expression of mRNA encoding ALK4 was reduced by 70% (Fig. 2F), with the remaining levels likely due to ALK4 expression in other EWAT cell types, including blood cells and resident immune cells, unaffected by the *Ap2*^CRE^ driver. Significant reductions were observed in the levels of mRNAs encoding Ki67, a marker of proliferating cells, and CD24 and ZFP423, two markers of adipocyte precursors, in EWAT of the mutant mice (Fig. 2F). While mutant EWAT showed a trend towards increased levels of mRNAs encoding PPARγ and CEBPα, two master regulators of adipogenic differentiation, they were not statistically different from those in EWAT of control mice (Fig. 2F). In order to assess the contribution of ALK4 activity towards adipose tissue mass in adult mice, we generated mice lacking ALK4 in adult adipose tissues by crossing *Acvr1b*^fx^ mice with *AdipoQ*^CRE-ERT2^ mice (22) and tamoxifen injection in *AdipoQ*^CRE-ERT2^::*Acvr1b*^fx/fx^ progeny at 1 month of age. We further stimulated fat mass expansion by subjecting mutant and control mice to HFD for 13 weeks beginning 2 weeks after the first tamoxifen injection. Expression of mRNA encoding ALK4 was reduced by 60% at the end of the experiment (Fig. 2G). However, the body weights of mutant and control mice were indistinguishable (Fig. 2H), as were the tissue weights of fat depots SWAT, EWAT and RWAT (Figs. 2I). Together, these results suggest that inactivation of ALK4 in adipose tissue from birth attenuates postnatal tissue expansion but has no effect on diet induced obesity in adult mice. Moreover, reduced expression of mRNAs encoding Ki67, CD24 and ZFP423 in adipose tissue of mice lacking ALK4 from birth suggests that ALK4 activity is required for the expansion of adipocyte precursors during neonatal adipose tissue development.

**Figure 2.**
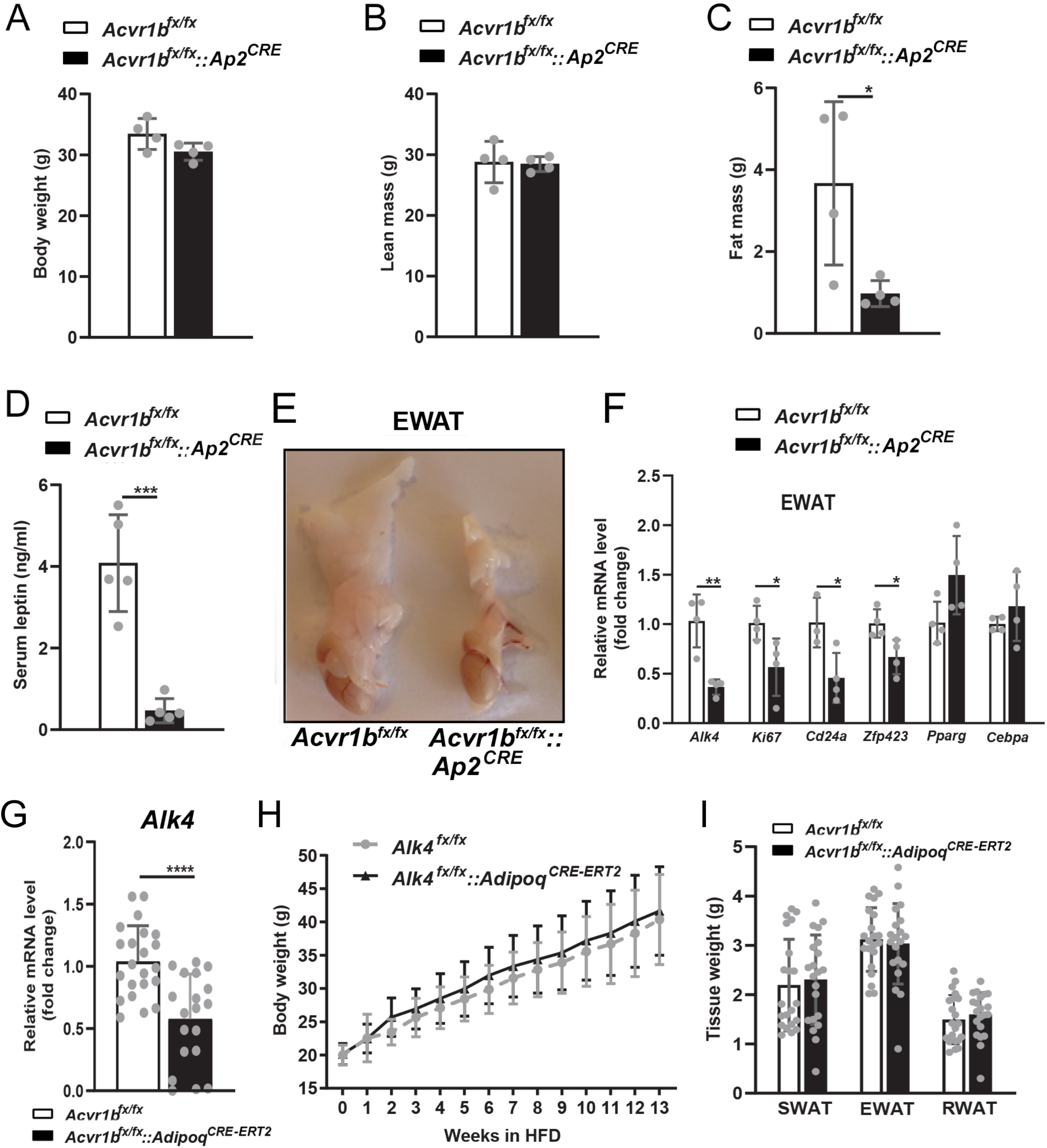
Inactivation of ALK4 in adipose tissue from birth attenuates postnatal tissue expansion but has no effect on diet induced obesity in adult mice. A-D) Body weight (A), lean mass (B), fat mass (C) and serum leptin levels in 4 month old control (*Acvr1b*^fx/fx^) and in mutant mice lacking ALK4 expression in adipose tissue from birth (*Acvr1b*^fx/fx^::*Ap2*^CRE^) kept under Chow diet. Data are presented as mean ± SD. *p<0.05 (unpaired t-test, N=5 mice per group). E) Macroscopic appearance of epididymal adipose tissue of 4 month old control (*Acvr1b*^fx/fx^) and mutant (*Acvr1b*^fx/fx^::*Ap2*^CRE^) mice kept under Chow diet. F) Expression of mRNAs encoding ALK4, Ki67, CD24a, Zfp423, PPARγ and CEBPα in EWAT of 4 month old control (*Acvr1b*^fx/fx^) and mutant (*Acvr1b*^fx/fx^::*Ap2*^CRE^) mice kept under Chow diet. mRNA levels were normalized to housekeeping genes Tbp and Ywhaz and plotted relative to control levels. Data are presented as mean ± SD. *p<0.05 ; *p<0.01 (2-way ANOVA, N=5 mice per group). G) Body weight of control (*Acvr1b*^fx/fx^) and mutant mice lacking ALK4 expression in adipose tissue from 1 month of age (*Acvr1b*^fx/fx^::*AdipoQ*^CRE-ERT2^) during high fat diet (HFD) feeding for 13 weeks (beginning 2 weeks after the first tamoxifen injection). Data are presented as mean ± SD. N=21 mice per group. H) Weights of dissected SWAT, EWAT and RWAT of control (*Acvr1b*^fx/fx^) and mutant (*Acvr1b*^fx/fx^::*AdipoQ*^CRE-ERT2^) mice after 13 week in high fat diet. Data are presented as mean ± SD. N=21 mice per group. I) Expression of mRNA encoding ALK4 in EWAT of control (*Acvr1b*^fx/fx^) and mutant (*Acvr1b*^fx/fx^::*AdipoQ*^CRE-ERT2^) mice after 13 week in HFD (N=8). mRNA levels were normalized to housekeeping genes Tbp and Ywhaz. Data are presented as mean ± SD. **p<0.01; (unpaired t-test).

### In contrast to ALK7, ALK4 is preferentially expressed in adipocyte precursors

Freshly isolated adipose tissues can be separated into a stromal vascular fraction (SVF), containing adipocyte precursors, macrophages, immune and endothelial cells, and a lighter fraction containing mature adipocytes. Expression of mRNA encoding ALK4 was enriched in SWAT and EWAT SVF isolated from of 6 week-old C57BL/6 mice (Fig. 3A). In contrast, Alk7 mRNA levels were very low or negligible in this fraction (Fig. 3A), suggesting predominant expression of ALK4, but not ALK7, in adipocyte precursors. In agreement with this, Alk4 mRNA expression declined in cultures of SWAT and EWAT SVF undergoing adipocyte differentiation (Fig. 3B). The initial increase in Alk4 mRNA expression observed in the early stage of EWAT SVF differentiation could be due to an additional round of proliferation of ALK4-expressing adipocyte precursors. In agreement with this, a similar pattern to that of Alk4 was observed in SWAT and EWAT for the adipocyte precursor marker Pdgfra (Fig. 1B). In contrast, Alk7 mRNA levels increased steadily, following the expression patterns of master regulators of adipogenesis PPARγ and CEBPα, as well as mRNAs encoding adipocyte markers adiponectin (AdipoQ) and fatty acid binding protein 4 (Fabp4, also known as Ap2) (Figs. 3B and C). Expression of mRNA encoding leptin, a marker of fully mature adipocytes, increased much later during differentiation of SVF cultures, as expected (Fig. 3C). Similar expression patterns were observed during differentiation of SVF isolated from human SWAT (Fig. 3D). In this case, the pattern of Alk4 and Pdgfra mRNA induction reSDbled that of mouse EWAT in that a transient increase was observed before their levels declined (Fig. 1D). SVF from mouse EWAT and human SWAT are more refractory to differentiation *in vitro* than mouse SWAT form the inguinal depot, suggesting a delayed cell cycle exit of adipocyte precursors of this tissues when placed under differentiation conditions. Together, these data indicate that ALK4 is primarily expressed in precursors of adipocytes and declines upon adipocyte differentiation in both subcutaneous and visceral mouse adipose tissues. In contrast, ALK7 is low or absent in adipocyte precursors and increases as differentiation proceeds, in parallel with other adipocyte markers such as adiponectin and Fabp4.

**Figure 3.**
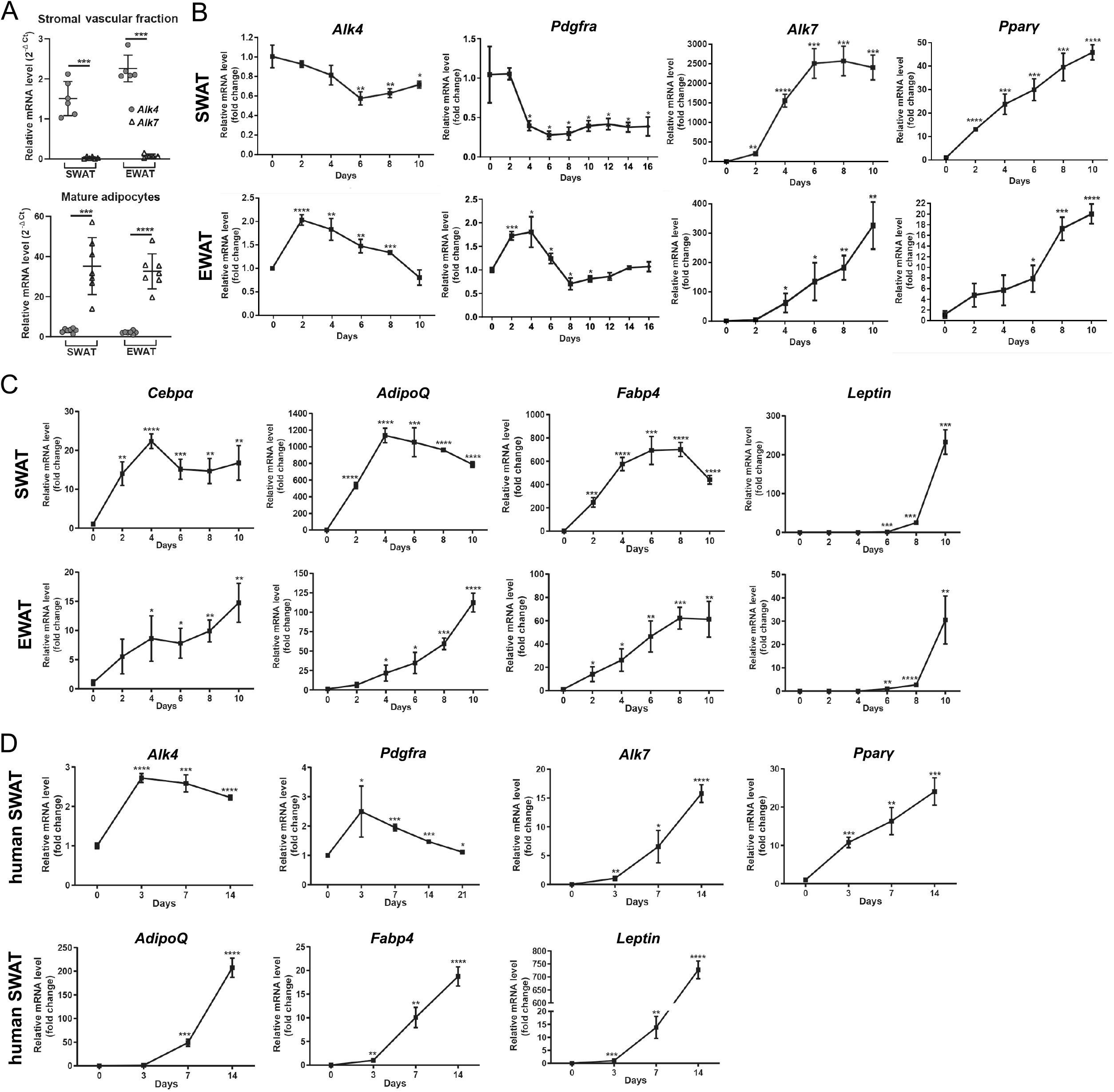
In contrast to ALK7, ALK4 is preferentially expressed in adipocyte precursors. A) Expression of mRNAs encoding ALK4 and ALK7 in SVF (upper graph) and mature adipocytes (lower graph) isolated from SWAT and EWAT of 6 week old mice. Each dot corresponds to one independent isolate. Relative mRNA levels were normalized to of housekeeping genes Tbp and Ywhaz. Means ± SD are indicated (N=6-7). ***p<0.001 (unpaired t-test). B) Expression of mRNAs encoding ALK4, Pdgfra, ALK7 and Pparγ during in vitro differentiation of SVF isolated from SWAT and EWAT of 6 week old mice. mRNA levels were normalized to housekeeping genes Tbp and Ywhaz and plotted relative to time 0. C) Similar to panel (B) for mRNAs encoding, Cebpα, adiponectin (AdipoQ), Fabp4 and Leptin. D) Expression of mRNAs encoding ALK4, Pdgfra, ALK7, Pparγ, adiponectin (AdipoQ), Fabp4 and Leptin during in vitro differentiation of SVF isolated from human SWAT biopsies. mRNA levels were normalized to housekeeping genes Tbp and Ywhaz and plotted relative to time 0. Data information: Data are presented as mean ± SD. *, p<0.05; **, p<0.01;***, p<0.001; ****, p<0.0001; unpaired t-test. N=3 biological replicates unless otherwise indicated.

### ALK4 signaling inhibits adipogenic differentiation and induces cell proliferation in SWAT and EWAT SVF cells

Expression of Alk4 mRNA in adipocyte precursors, together with lipodystrophy in mutant mice lacking ALK4 in adipose tissue, suggested a role for ALK4 in the differentiation and proliferation of pre-adipocytes. We tested this notion in SVF cultures subjected to adipogenic differentiation. Activation of ALK4 signaling by treatment with activin A during SVF differentiation dramatically reduced Oil Red O staining of lipids in adipocytes derived from SWAT and EWAT SVF (Fig. 4A), indicating that ALK4 signaling inhibits adipogenic differentiation of these cells. The effects of activin A on cell proliferation was assessed during early stages of SVF differentiation by exposing 2 day -old activin A-treated cultures to BrdU during the last 30min. In both SWAT and EWAT cultures did activin A enhance BrdU incorporation compared to vehicle treatment (Fig. 4B), indicating that reduced adipogenic differentiation by ALK4 signaling is paralleled by increased proliferation of adipocyte precursors.

**Figure 4.**
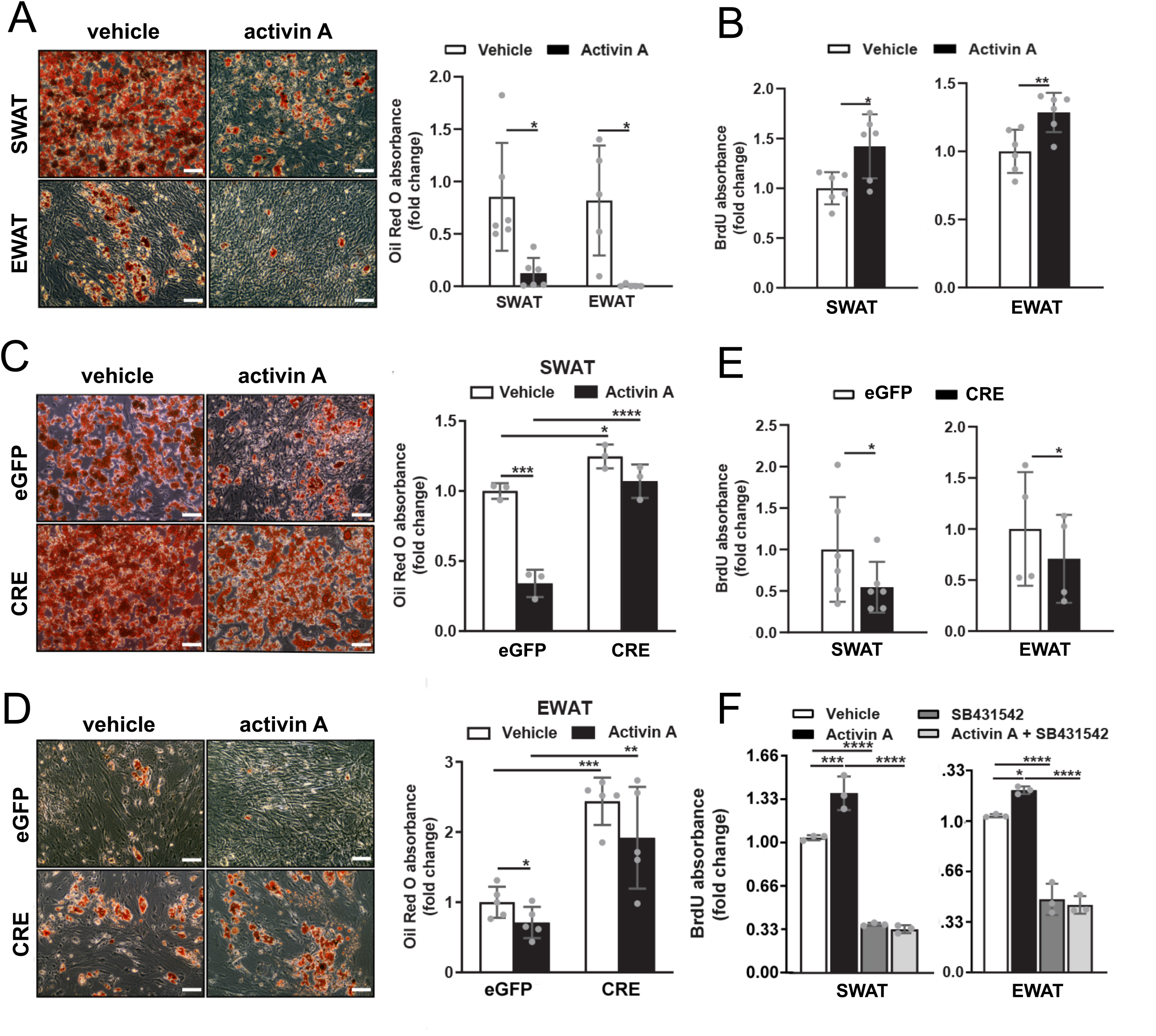
ALK4 signaling inhibits adipogenic differentiation and induces cell proliferation in SWAT and EWAT SVF cells. A) Wild type SVF of 6 week old mouse SWAT and EWAT were induced to differentiate in vitro during treatment with vehicle or activin A for 8 days, then stained with Oil Red O. Histogram shows fold change Oil red O absorbance relative to vehicle. B) Wild type SVF of 6 week old mouse SWAT and EWAT were induced to differentiate in vitro during treatment with vehicle or activin A for 2 days, then assessed for BrdU incorporation during the last 30 min of culture. Histogram shows fold change BrdU absorbance relative to vehicle. C) SVF of 6 week old *Acvr1b*^fx/fx^ mouse SWAT were induced to differentiate for 8 days in vitro after infection with control (eGFP) or CRE-expressing (CRE) adenoviruses, treated with vehicle or activin A as indicated, then stained with Oil Red O. Histogram shows fold change Oil red O absorbance relative to eGFP samples treated with vehicle. D) Similar to panel (C) for EWAT. E) SVF of 6 week old *Acvr1b*^fx/fx^ mouse SWAT and EWAT were induced to differentiate for 2 days in vitro after infection with control (eGFP) or CRE-expressing (CRE) adenoviruses, treated with vehicle or activin A as indicated, then assessed for BrdU incorporation during the last 30 min of culture. Histogram shows fold change BrdU absorbance relative to eGFP samples treated with vehicle. F) Wild type SVF of 6 week old mouse SWAT and EWAT were induced to differentiate in vitro during treatment with vehicle (white) activin A (black), SB431542 ALK4 kinase inhibitor (dark grey) or activin A plus SB431542 (light grey) for 8 days, then assessed for BrdU incorporation during the last 30 min of culture. Histogram shows fold change BrdU absorbance relative to vehicle. Data information: Data are presented as mean ± SD. *, p<0.05; **, p<0.01;***, p<0.001; ****, p<0.0001; one-way ANOVA followed with Tukey’s Sidak’s multiple comparison test. N=3 biological replicates.

Inactivation of ALK4 was achieved using CRE-expressing adenoviruses in cultures of SVF isolated from SWAT and EWAT of 6 week-old *Acvr1b*^fx/fx^ mice. Oil Red O staining was increased in both SWAT and EWAT SVF cultures after infection with CRE-expressing viruses compared to eGFP-expressing controls (Figs. 4C and D). Treatment with activin A reduced Oil Red O staining in control cultures but had no significant effects in CRE-expressing cultures, confirming the requirement of ALK4 for activin A signaling (Figs. 4C and D). In SVF cultures from *Acvr1b*^fx/fx^ SWAT and EWAT, infection with CRE-expressing adenovirus decreased BrdU incorporation compared to eGFP control virus infection (Fig. 4E), in opposition to the effects of activin A stimulation. Finally, we tested the effects of the ALK4 kinase inhibitor SB431542 on BrdU incorporation in SWAT and EWAT SVF cultures treated with activin A. Again activin A enhanced BrdU incorporation, but this effect was blocked by the SB431542 inhibitor (Fig. 4F). Interestingly, the inhibitor was also able to suppress BrdU incorporation on its own, compared to vehicle-treated cells (Fig. 4F), indicating the presence of endogenous ALK4 ligands in the cultures. In conclusion, ligand activation as well as genetic or pharmacological inactivation of ALK4 consistently affect pre-adipocyte differentiation and proliferation in a converse fashion, indicating that ALK4 functions is to dampen differentiation of adipocyte precursors to allow their proliferation, thus promoting adipose tissue expansion through hyperplasia.

### ALK4 signaling suppresses expression of adipogenic genes

In order to get a better understanding of the effects of ALK4 activation and inactivation in adipogenic differentiation, we subjected SVF cultures from SWAT and EWAT of 6 week-old C57BL/6 mice to differentiation in the presence or absence of activin A and assessed the expression dynamics of key adipogenic marker genes through the differentiation process. The same set of markers was also investigated in cultures of SVF isolated from SWAT and EWAT of 6 week old *Acvr1b*^fx/fx^ mice after infection with CRE-expressing or eGFP-expressing adenoviruses. Treatment with activin A transiently increased mRNA encoding ALK4 in SWAT, but not in EWAT (Fig. 5A), perhaps reflecting a transient amplification of adipocyte precursors in the early stages of the culture. On the other hand, activin A strongly repressed expression of all marker genes of differentiated and mature adipocytes, including Alk7, Pparγ, Cebpα, AdipoQ, Fabp4 and leptin, in both SWAT and EWAT (Fig. 5A). Conversely, infection with CRE-expressing adenovirus of *Acvr1b*^fx/fx^ SVF cultures significantly enhanced the expression of these genes (Fig. 5B). Interestingly, activin A treatment increased, but Alk4 deletion decreased, expression of mRNA encoding CD24, a marker of adipocyte progenitors (23) (Fig. 5C), in agreement with the role of ALK4 signaling in mitotic-competent adipocyte precursors. Importantly, activin A also suppressed expression of the same set of marker genes during differentiation of SVF isolated from human SWAT (Fig. 5D). Thus, ALK4 activation suppresses, while ALK4 inactivation enhances, adipocyte differentiation by regulating the expression of genes responsible for the induction of the mature adipocyte phenotype and the maintenance of the precursor state.

**Figure 5.**
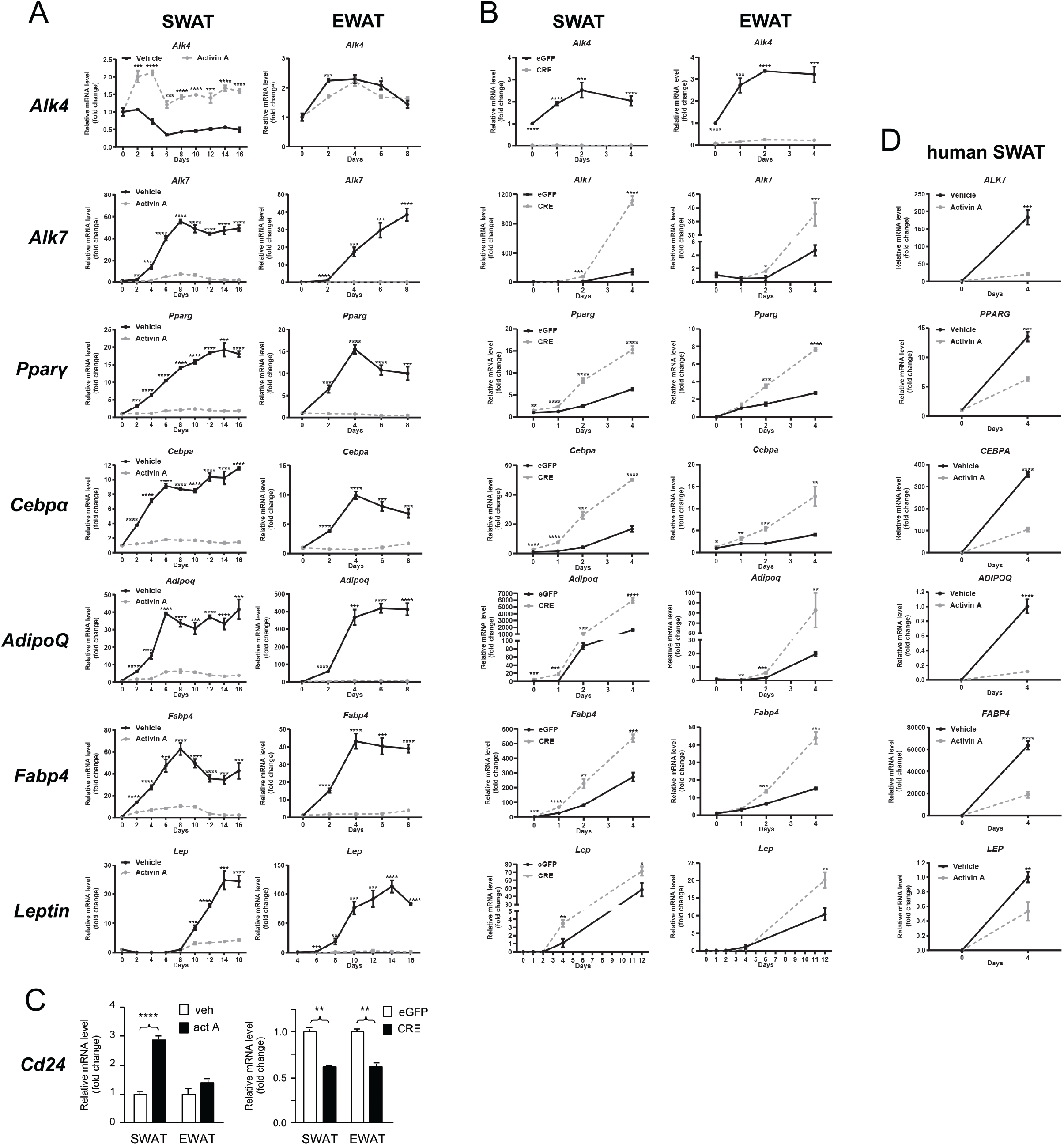
ALK4 signaling suppresses expression of adipogenic genes. A) Expression of mRNAs encoding ALK4, ALK7, Pparγ, Cebpα, adiponectin (AdipoQ), Fabp4 and Leptin during in vitro differentiation of SVF isolated from SWAT and EWAT of 6 week old mice in the presence (grey dotted lines) or absence (black sold lines) of activin A. mRNA levels were normalized to housekeeping genes Tbp and Ywhaz and plotted relative to time 0. B) Expression of mRNAs encoding ALK4, ALK7, Pparγ, Cebpα, adiponectin (AdipoQ), Fabp4 and Leptin during in vitro differentiation of SVF isolated from and SWAT and EWAT of 6 week old *Acvr1b*^fx/fx^ mice after infection with control (eGFP, black solid lines) or CRE-expressing (CRE, grey dotted lines) adenoviruses. Gene labels as in the left hand side. mRNA levels were normalized to housekeeping genes Tbp and Ywhaz and plotted relative to time 0. C) Expression of mRNA encoding CD24 after 2 day in vitro differentiation of SVF isolated from SWAT and EWAT of 6 week old mice in the presence (white bar) or absence (black bar) of activin A (upper panel), or SVF from *Acvr1b*^fx/fx^ mice after infection with control (eGFP, black solid lines) or CRE-expressing (CRE, grey dotted lines) adenoviruses. mRNA levels were normalized to housekeeping genes Tbp and Ywhaz and plotted relative to control conditions. D) Expression of mRNAs encoding ALK7, Pparγ, Cebpα, adiponectin (AdipoQ), Fabp4 and Leptin during in vitro differentiation of SVF isolated from human SWAT biopsies in the presence (grey dotted lines) or absence (black sold lines) of activin A. Gene labels as in the left hand side. mRNA levels were normalized to housekeeping genes Tbp and Ywhaz and plotted relative to time 0. Data information: Data are presented as mean ± SD. *, p<0.05; **, p<0.01;***, p<0.001; ****, p<0.0001; unpaired t-test. N=3 biological replicates unless otherwise indicated.

### ALK4 signaling preferentially activates Smad2 and downregulates the levels of PPARγ and CEBPα proteins during adipogenic differentiation

We examined the phosphorylation status of Smad2 and Smad3, the two main immediate downstream effectors of ALK4 signaling in cultures of differentiated SWAT and EWAT SVF from *Acvr1b*^fx/fx^ mice treated with CRE virus or eGFP control after stimulation with activin A or vehicle. ALK4 activation by activin A induced mainly (SWAT, Fig. 6A) or exclusively (EWAT, Fig. 6B) phosphorylation of Smad2, which was abrogated by infection with CRE-expressing adenovirus. The expression of PPARγ and CEBPα proteins was reduced by treatment with activin A, but significantly increased after CRE-mediated Alk4 deletion with adenovirus (Figs. 6A and B). Together, these indicate that ALK4 signaling preferentially activates Smad2 and downregulates the levels of PPARγ and CEBPα proteins during adipogenic differentiation of adipocyte precursors.

**Figure 6.**
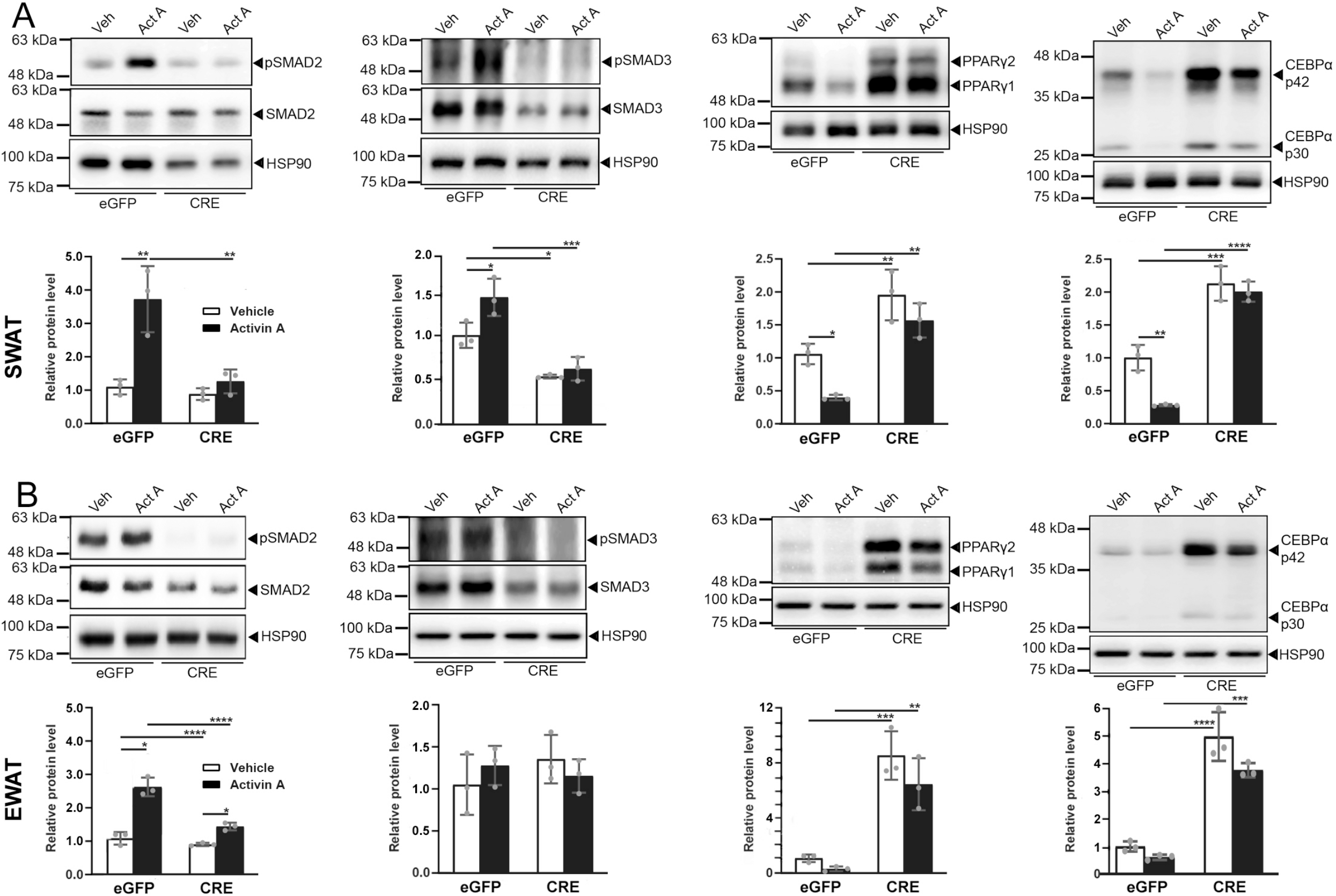
ALK4 signaling induces preferential phosphorylation of Smad2 and decreases levels of PPARγ and CEBPα proteins during adipogenic differentiation. A) Phosphorylation of Smad2 and Smad3 and protein expression of PPARγ and CEBPα after 3 day differentiation of SVF isolated from SWAT of 6 week old *Acvr1b*^fx/fx^ mice after infection with control (eGFP) or CRE-expressing adenoviruses in the presence (act A) or absence (ver) of activin A. Histograms (white bars vehicle, black bars activin A) show fold change relative to vehicle/eGFP normalized to total Smad2 (for pSmad2), total Smad3 (for pSmad3) or HSP90 (for PPARγ and CEBPα), respectively. B) Similar to panel (A) for EWAT. Data information: Data are presented as mean ± SD. *, p<0.05; **, p<0.01;***, p<0.001; ****, p<0.0001; unpaired t-test. N=3 biological replicates.

### Knock-down of *Cebpα* counteracts the effects of *Alk4* deletion in adipogenic differentiation

In order to confirm the role of CEBPα in the ability of ALK4 signaling to suppress adipogenic differentiation, we tested whether diminishing CEBPα levels in preadipocytes lacking ALK4 would restore their differentiation to normal levels. SWAT and EWAT SVF from *Acvr1b*^fx/fx^ mice were infected with either eGFP or CRE-expressing adenoviruses, simultaneously transfected with siRNA targeting Cebpα mRNA or a control (scrambled) siRNA, and then subjected to adipocyte differentiation. Confirming our previous results, inactivation of ALK4 with CRE adenoviruses enhanced the expression of mRNA encoding adipogenic markers Pparγ and Cebpα (Fig. 7A), as well as that of mRNAs encoding markers of more mature adipocytes, such as ALK7, adiponectin (AdipoQ), Fabp4 and leptin (Fig. 7B). siRNA knock-down of Cebpα in pre-adipocytes reduced its mRNA levels by 50-60% (Fig. 7A). Importantly, in adipocyte precursors lacking ALK4, reducing the levels of Cebpα largely restored adipogenic differentiation to near normal levels, as the expression of mRNA encoding adipogenic and maturation markers returned to levels similar to those of cells that received control adenovirus and siRNA (Figs. 7A and B). Interestingly, the expression levels of mRNA encoding ALK4 increased after Cebpα knock-down (Fig. 7A), suggesting a reciprocal negative regulation between these two proteins.

**Figure 7.**
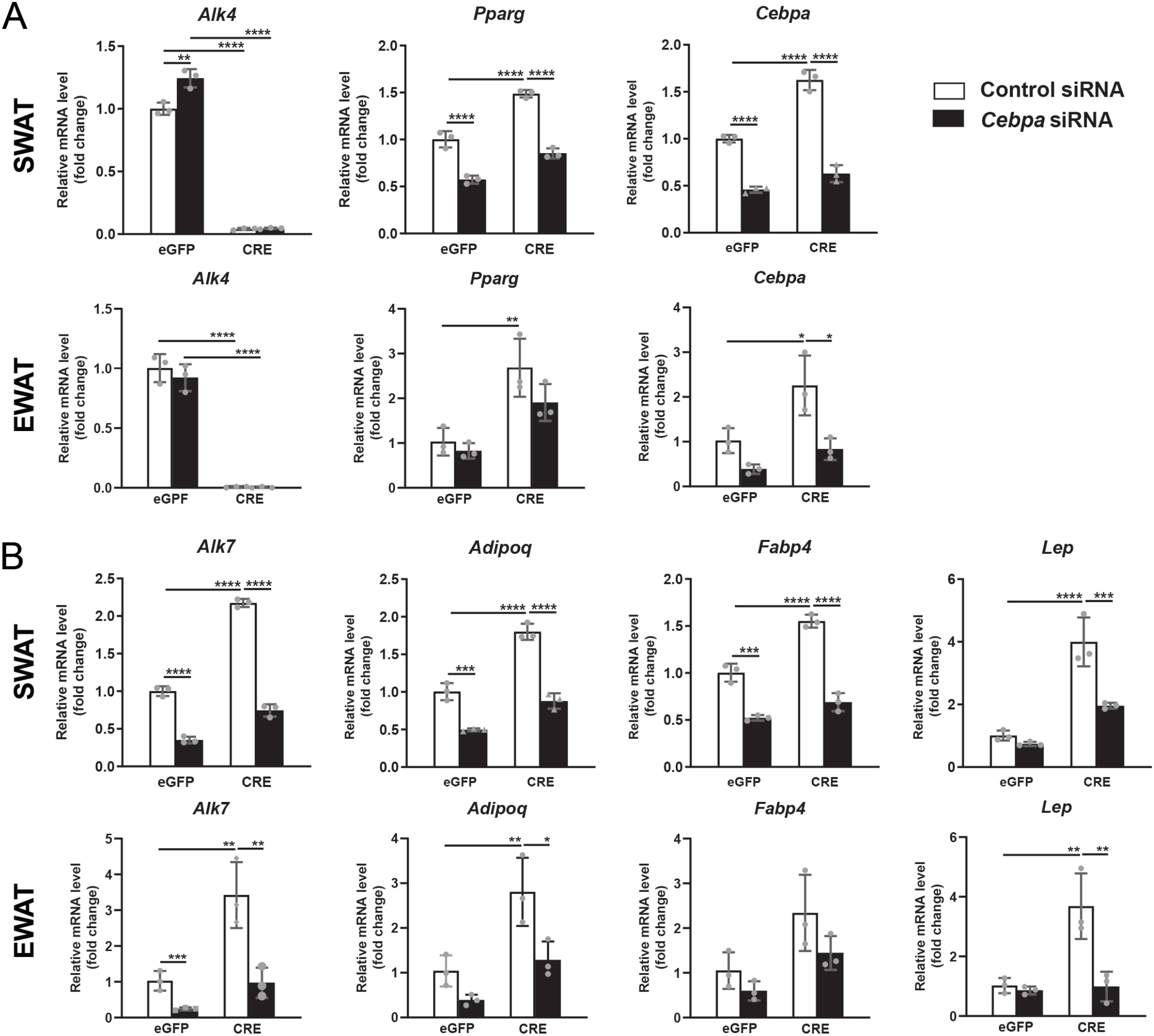
Knock-down of *Cebpα* counteracts the effects of *Alk4* deletion in adipogenic differentiation. A) Expression of mRNAs encoding ALK4, Pparγ and Cebpα, after 4 day differentiation of SVF isolated from SWAT or EWAT of 6 week old *Acvr1b*^fx/fx^ mice after infection with control (eGFP) or CRE-expressing adenoviruses and transfection with control siRNA (white bars) or Cebpα siRNA (black bars) as indicated. mRNA levels were normalized to housekeeping genes Tbp and Ywhaz and are presented as fold change relative to eGFP/control siRNA. B) Similar to panel (A) for mRNAs encoding ALK7, adiponectin (AdipoQ), Fabp4 and Leptin. Data information: Data are presented as mean ± SD. *, p<0.05; **, p<0.01;***, p<0.001; ****, p<0.0001; unpaired t-test. N=3 biological replicates.

## Discussion

In this study, we examined the expression dynamics, regulation and function of the canonical activin receptor ALK4 in subcutaneous and visceral adipose tissues during low and high caloric regimes. The ALK4 protein sequence is only 30% identical to that of ALK7 in the extracellular domain, which explains their differential ability to interact with TGF-β superfamily ligands. In their intracellular domains, however, the two proteins are over 80% identical, and so their signaling capabilities are remarkably close (17). One of the main findings of this study is the contrasting functions of these receptors. We found that ALK4 impacts adipose tissue hyperplasia by dampening the differentiation, thus enhancing the proliferative capacity, of adipocyte precursors. On the other hand, ALK7 affects adipose tissue hypertrophy by suppressing catabolic pathways, eg lipolysis and beta-oxidation, in mature adipocytes (10). In adipose tissue, the two main ligands of these receptors are activin B and GDF-3 (reviewed in (17)). Although activin A can only interact with ALK4, the levels of this ligand are low in adipose tissue and largely unaffected by caloric intake (this study and (9)). Therefore, given the high similarities in intracellular protein sequence and signaling capabilities between ALK4 and ALK7, we attribute their disparate regulation and function to their differential expression during early, ALK4, and later, ALK7, stages of adipogenic differentiation, respectively.

In a low caloric regime, mammals balance their energy consumption by reducing their metabolic rate, resulting is accelerated weight gain when subsequently placed *ad libitum* on a high caloric diet (24, 25). A recent study found that pair-feeding mice that underwent diet-induced weight reduction with lean control animals efficiently stabilized body weight, indicating that hyperphagia may also be an important contributor to weight regain after a diet change (26). In line with this, we found that the caloric intake on HFD of mice undergoing diet switch consistently overshot that of animals placed under constant HFD. Accordingly, the weights of visceral adipose tissues, EWAT and RWAT, after the switch group was placed on HFD surpassed the projected tissue weights observed in mice under constant HFD. We propose that downregulation of ALK4 together with concomitant upregulation of ALK7 promotes differentiation and hypertrophy of adipocytes during the low caloric phase of the diet cycle. After a return to HFD, ALK4 upregulation promotes further expansion of the tissue by hyperplasia. This could explain how repeated cycles of low and high caloric intake can result in adipose tissue expansion that can go beyond that observed under a constant high caloric regime.

Although ALK4 and ALK7 affect adipogenic differentiation in opposite directions, they both contribute to tissue expansion by their respective effects on hyperplasia and hypertrophy. In fact, the lipodystrophic phenotype of *Ap2*^CRE^::*Acvr1b*^fx/fx^ mutant mice (lacking ALK4) kept in Chow diet (this study) was even more dramatic than that of *Ap2*^CRE^::*Acvr1c*^fx/fx^ mice (lacking ALK7) under the same diet regime (10). We attribute the drastically reduced fat mass of the ALK4 mutants to premature cell cycle exit and differentiation of adipocyte precursors, preventing appropriate tissue expansion during postnatal development. In line with this, inactivation of ALK4 during adult stages had no effect on total body weight or adipose tissue mass under constant high fat diet. Future studies should address the consequence of adult ALK4 inactivation on animals subjected to repeated cycles of high a low caloric diets.

*In vitro* studies of cultured SVF allowed us to follow longitudinally the proliferation, differentiation and maturation of a synchronized population of cells during the course of adipogenesis as well as test the effects of loss and gain of ALK4 function in this process. Activation of ALK4 by stimulation with activin A dramatically suppressed adipocyte differentiation as revealed by aborted induction of adipogenic master genes Pparγ and Cebpα, and later markers of differentiated and mature adipocytes, including ALK7, adiponectin, Fabp4 and leptin. Since activin A is unable to activate ALK7, we are confident that these effects are mediated by the ALK4 receptor. The fact that activin A had similar effects in cells derived from human SVF indicate a conserved function of ALK4 in human fat. In contrast, fat-specific inactivation of ALK4 by CRE-mediated deletion accelerated adipogenesis as revealed by premature upregulation of all adipogenic and maturation markers in both subcutaneous and visceral adipocytes.

A previous study in pancreatic islets reported that activins A and B induce differential phosphorylation of Smad2 and Smad3, respectively, suggesting that ALK4, the main activin A receptor, may signal preferentially through Smad2 over Smad3 in these cells (27). Other studies showed that activin A activates the Smad2 pathway in human adipocyte progenitors (7) and in the 3T3-L1 preadipocyte cell line (28). In agreement with those observations, we found that activation of ALK4 by activin A induced Smad2 phosphorylation to a greater extent than Smad3 in cultured SWAT and EWAT SVF cells. Activin A decreased the levels of PPRAγ and CEBPα proteins, two indispensable drivers of adipogenic differentiation, while CRE-mediated inactivation of ALK4 increased their levels by 5 to 10 fold, respectively, indicating increased differentiation potential in adipocyte precursors lacking ALK4. That the effects of ALK4 on adipogenesis are mediated by suppression of CEBPα expression was corroborated by the restoration of close to normal levels of adipogenic differentiation after Cebpα knock-down in SVF cultures derived from ALK4 mutant mice. Earlier studies indicated that Smad3 can directly interact with CEBPβ and CEBPδ, two key upstream factors necessary for induction of CEBPα expression, and thereby suppress their function (5). Although Smad2 was not found to interact with CEBP proteins, those studies, performed using *in vitro* translated, unphosphorylated Smad proteins, did not address the possible involvement of endogenous Smad2 in primary adipocytes after phosphorylation by the ALK4 kinase. Interestingly, another study found that knockdown of both Smad2 and Smad3 was required to elevate the levels of PPARγ and CEBPα proteins in the adipocyte cell line 3T3-L1 (11), suggesting that Smad2 may be necessary but not sufficient to regulate PPARγ and CEBPα levels. It is therefore possible that Smad2 interacts indirectly with adipogenic factors as part of a Smad2/3 complex and additional studies will be required to address this possibility.

In conclusion, the results of the present study reveal an intriguing contrast between the expression patterns, dynamics and function of the two main type I activin receptors ALK4 and ALK7. While the former promotes hyperplasia by delaying differentiation of adipocyte precursors (this study), the latter promotes hypertrophy by suppressing catabolic pathways, such as lipolysis and beta-oxidation (10). Through the promotion of sequential cycles of hyperplasia followed by hypertrophy, the combined action of these two receptors may be responsible for enhanced adipose tissue expansion during repeated cycles of high and low caloric intake.

## Experimental Procedures

### Mice

C57BL/6NTac mice were kept on a 12 hour light-dark cycle with ad libitum access to standard chow diet (irradiated 2018 Teklad Global Rodent Diet; Envigo, Madison, USA) and water. *Acvr1b*^fx/fx^ conditional knockout mice as previously described (20)) were also used in the experiment. Adipose tissue-specific knock-out lines, *Acvr1b*^fx/fx^::*Ap2*^Cre^ and *Acvr1b*^fx/fx^::*Adipoq*^Cre^ were generated by crossing Acvr1bfx/fx mice with *Ap2*^Cre^ (The Jackson Laboratory, USA, 018965) and *Adipoq*^Cre^ mice (22), respectively. To induce cre recombinase-mediated *Acvr1b* deletion, 4-week-old *Acvr1b*^fx/fx^ and *Acvr1b*^fx/fx^:: *Adipoq*^Cre^ mice were injected intraperitoneally with tamoxifen (Sigma-Aldrich; T5648) prepared in saline solution containing 5% DMSO and 5% Tween-80, at a concentration of 100 μg/10 g body weight for 5 consecutive days. Only male mice were used for the study. All animal procedures were approved by the National University of Singapore Institutional Animal Care and Use Committee and by the Stockholms Norra Djurförsöksetiska Nämnd.

### High fat diet and dietary switch

Male mice at 6 weeks of age were fed with either a standard chow diet, comprises 5% kcal from fat (irradiated 2018 Teklad Global Rodent Diet; Envigo) or a high fat diet, comprises 60% kcal from fat (Research Diets, New Brunswick, USA) for 13 or 14 weeks. Body weight was measured every week. Adipose tissues were collected from inguinal subcutaneous (SWAT), epididymal (EWAT) and retroperitoneal (RWAT) depots, weighted, washed twice with Hank’s balanced salt solution (Nacalai Tesque Inc, Japan, 17461-05) and subjected to RNA isolation. For the dietary switch study, male mice at 6 weeks of age were fed with 4 weeks of high fat diet. Thereafter, the mice were divided into 2 groups and were subjected to different nutritional interventions for 10 weeks. The control group was fed with high fat diet constantly and the experimental group was subjected to diet switch in this chronological order: chow diet (2 weeks) – high fat diet (2 weeks) –chow diet (2 weeks) – high fat diet (4 weeks). Body weight and food consumption were measured every week. Mice were sacrificed for collection of adipose tissues upon switch of diet. Adipose tissues were collected from inguinal subcutaneous, epididymal and retroperitoneal depots, weighted, washed twice with Hank’s balanced salt solution and subjected to histological analysis, stromal vascular fraction isolation and RNA isolation.

### Mouse and human SVF isolation, culture and adipogenic differentiation

Male C57BL/6NTac mice at 6 weeks of age were sacrificed for isolation of SVF. Adipose tissues were isolated from inguinal subcutaneous and epididymal depots, washed twice with Hank’s balanced salt solution and minced with scissors. Minced tissue was digested for 60 min with 1 mg/ml collagenase type I (Sigma-Aldrich, USA, C0130) in digestion buffer, composed of, 0.1 M N-2-hydroxyethylpiperazine-N’-2-ethanesulfonic acid (HEPES) sodium salt (Sigma-Aldrich, H7006), 0.12 M sodium chloride (Sigma-Aldrich), 50 mM potassium chloride (Sigma-Aldrich), 5 mM d-glucose (Sigma-Aldrich), 1 mM calcium chloride (Sigma-Aldrich, Japan) and 1.5% bovine serum albumin (BSA, Biowest, France) with constant shaking. Digested tissue was filtered through a 100 μm cell strainer, then through a 20 μm nylon mesh. Filtrate was centrifuged for 5 min at 100 rcf to separate floating mature adipocytes. After removing mature adipocytes, supernatant was further centrifuged for 6 min at 250 rcf. The resulting cell pellet was washed with basal medium (advanced DMEM/F12, Gibco, USA, 12634028), containing 5% fetal bovine serum (Gibco, 10500) and 100 U/ml penicillin-streptomycin (Gibco, 15140122). Cells were repelleted, plated in cell culture dish and grew in advanced DMEM/F12, supplemented with 10% heat inactivated fetal bovine serum, 2 mM Glutamax (Gibco, 35050061), 0.1 mM non-essential amino acids (Gibco, 11140050) and 100 U/ml penicillin-streptomycin. After overnight adherence, cells were gently washed with basal medium and expanded for experiments. All cell cultures were performed in an incubator at 37°C with 5% CO2. SVF at passage 1 to 3 were used for adipogenic differentiation. Cells were seeded on 0.1% gelatin (Sigma-Aldrich, G9391) coated plates and cultured to confluence. Two days after confluence, cells were subjected to adipogenic differentiation in advanced DMEM/F12 supplemented with 5% heat inactivated fetal bovine serum, 33 μM d-biotin (Sigma-Aldrich, B4639), 17 μM pantothenate (Sigma-Aldrich, P5155), 0.5 mM 3-isobutyl-1-methylxanthine (Sigma-Aldrich, I5879), 1 μM dexamethasone (Sigma-Aldrich, D4902), 1 μM rosiglitazone (Sigma-Aldrich, R2408), 2 nM 3,3′,5-Triiodo-L-thyronine sodium salt (Sigma-Aldrich, T6397), 1 ug/ml insulin, 0.55 μg/ml transferrin, 0.67 ng/ml sodium selenite (Gibco, 41400045), 2 mM Glutamax, 0.1 mM non-essential amino acids and 100 U/ml penicillin-streptomycin for 4 days. Differentiated cells were maintained in advanced DMEM/F12 supplemented with 2% heat inactivated fetal bovine serum, 33 μM d-biotin, 17 μM pantothenate, 1 μM dexamethasone, 10 ug/ml insulin, 2 mM Glutamax, 0.1 mM non-essential amino acids and 100 U/ml penicillin-streptomycin until full differentiation (i.e. day 8 – 16). Medium was changed every 2 days in the period of differentiation and maintenance.

For isolation of human SVF, white adipose tissues were dissected from abdominal subcutaneous depot of human subjects who underwent bariatric surgery. Procedures for isolation and culture of mouse SVF were adopted. All procedures are in accordance with guidelines of the National Healthcare Group Domain Specific Review Board, Singapore. Human stromal vascular fractions at passages 1 to 3 were used for adipogenic differentiation. Cells were seeded on gelatin coated plates and cultured to confluence. Two days after confluence, cells were subjected to adipogenic differentiation in advanced DMEM/F12 supplemented with 5% heat inactivated fetal bovine serum, 33 μM d-biotin, 17 μM pantothenate, 0.5 mM 3-isobutyl-1-methylxanthine, 1 μM dexamethasone, 1 μM rosiglitazone, 2 nM triiodothyronine, 1 ug/ml insulin, 0.55 μg/ml transferrin, 0.67 ng/ml sodium selenite, 2 mM Glutamax (Gibco), 0.1 mM non-essential amino acids (Gibco) and 100 U/ml penicillin-streptomycin for 7 days. Differentiated cells were maintained in advanced DMEM/F12 supplemented with 2% heat inactivated fetal bovine serum, 33 μM d-biotin, 17 μM pantothenate, 1 μM dexamethasone, 10 ug/ml insulin, 2 mM Glutamax (Gibco), 0.1 mM non-essential amino acids (Gibco) and 100 U/ml penicillin-streptomycin until full differentiation (i.e. day 14). Medium was changed every 2 days in the period of differentiation and maintenance.

### CRE-mediated ALK4 inactivation in *Acvr1b*^fx/fx^ mouse SVF

SVFs were isolated from inguinal subcutaneous and epididymal depots of 6-week-old male *Acvr1b*^fx/fx^ mice and cultured as described earlier. All adenoviruses used are replication defective human adenovirus type 5 (E1/E3 deleted) having cyto-megalovirus (CMV) promoter and enhanced green fluorescent protein (eGFP) as reporter. A total of 45,000 cells were mixed with either control eGFP adenovirus (Vector Biolabs, USA, 1060) that encodes eGFP or cre recombinase-eGFP adenovirus that encodes cre recombinase and eGFP in tandem (Vector Biolabs, 1700) at 400 multiplicity of infection and 6 μg/ml polybrene (Sigma-Aldrich, H9268) before being seeded on 0.1% gelatin-coated 24-well plate in advanced DMEM/F12, supplemented with 10% heat inactivated fetal bovine serum, 2 mM Glutamax, 0.1 mM non-essential amino acids and 100 U/ml penicillin-streptomycin. After 48 h, culture medium was refreshed and infected cells were used for experiments on the following day.

### siRNA-mediated knock-down of Cebpα gene expression in *Acvr1b*^fx/fx^ mouse SVF

Modified reverse siRNA transfection of earlier published protocol (Isidor et al, 2016) was adopted. Briefly, Lipofectamine 2000 Reagent (Invitrogen, USA, 11668027) and 20 μM siRNAs targeting Cebpa, ID: s63854 and s63855 (Ambion, USA, 4390771) were diluted separately in Opti-MEM Reduced Serum Medium (Gibco, 51985034). Silencer Select Negative Control #1 siRNA (Ambion, 4390843) was used as scrambled control. Equal volume of diluted Lipofectamine and siRNAs were mixed and transferred to 0.1% gelatin-coated 24-well plate. Lipofectamine and siRNA mixtures at final concentration of 10 μl/ml and 25 nM, respectively were incubated for 25 min at room temperature. SVF from inguinal subcutaneous and epididymal depots of 6-week-old male Acvr1bfx/fx mice were mixed with either control eGFP adenovirus or cre recombinase-eGFP adenovirus at 400 multiplicity of infection and 6 μg/ml polybrene in advanced DMEM/F12, supplemented with 10% heat inactivated fetal bovine serum, 2 mM Glutamax and 0.1 mM non-essential amino. Cells (45,000 cells per well) and viruses mixture were transferred to 24-well plate preincubated with lipofectamine and siRNA mixture. After 48 h, culture medium was refreshed and cells were subjected to adipocyte differentiation on the following day.

### Bromodeoxyuridine (BrdU) incorporation assay

SVF were seeded at a density of 30 000 cells/cm2 on 0.1% gelatin-coated plates and left for 48 h before subjected to adipocyte differentiation. For examination of cell proliferation, cells were stimulated with 100 ng/ml activin A and/ or 10 μM SB431542 (Sigma-Aldrich, S4317) in advanced DMEM/F12 supplemented with 5% heat inactivated fetal bovine serum, 33 μM d-biotin, 17 μM pantothenate, 0.5 mM 3-isobutyl-1-methylxanthine, 1 μM dexamethasone, 1 μM rosiglitazone, 2 nM triiodothyronine, 1 ug/ml insulin, 0.55 μg/ml transferrin, 0.67 ng/ml sodium selenite, 2 mM Glutamax (Gibco), 0.1 mM non-essential amino acids (Gibco) and 100 U/ml penicillin-streptomycin for 12 h (SWAT) or 24 h (EWAT). Stimulated cells were incubated with 10 μM BrdU labeling reagent for 30 min, washed with PBS, fixed and proceeded with colorimetric immunoassay for the quantification of cell proliferation according to manufacturer instructions (Roche, Germany, 11647229001).

### Oil Red O staining of differentiated adipocyte cultures

On day 8 of SVF differentiation, cells were washed with PBS, fixed with 4% paraformaldehyde for 10 min and subjected to Oil red O staining. Oil Red O (Sigma-Aldrich, O0625) stock solution, 3.5 mg/ml was prepared in absolute isopropanol. Oil Red O working solution was prepared by mixing 3 parts of Oil Red O stock solution with 2 parts of deionized water. Before using, the working solution was left for 10 min at room temperature and filtered through 0.2 μm filter. Fixed cells were washed with 60% isopropanol and stained with Oil Red O working solution for 1 h at room temperature. Stained cells were washed with deionized water and 60% isopropanol. To quantify staining, Oil Red O dye was eluted with absolute isopropanol, comprises 4% NP-40. Optical density (OD) was measured at a wavelength of 500 nm and normalized to total DNA content. For quantification of DNA content, cells were washed with absolute isopropanol and 75% ethanol. Cells were then air-dried for 30 min and lyse with 200 μg/ml proteinase K (Promega, WI, USA) in lysis buffer, comprises 50 mM potassium chloride (Sigma-Aldrich), 10mM Tris-HCL (pH 9; 1st Base, Singapore) and 0.1% Triton X-100 (Sigma-Aldrich) at 60°C for 1 h. Amount of DNA was quantified with NanoDrop 2000 spectrophotometer (ThermoFisher Scientific, USA). Images of stained cells were acquired using an inverted microscope (Zeiss AX10, Germany) and a charged-coupled device (CCD) camera (QImaging Retiga R6, USA) with Ocular Imaging software (QImaging).

### RNA isolation and quantitative RT-PCR

Adipose tissues or cells were homogenized in TRIzol reagent (Invitrogen). RNA was isolated according to the manufacturer instructions. RNA concentration and purity were assessed with NanoDrop 2000 spectrophotometer (ThermoFisher Scientific). Total RNA was reverse-transcribed using the FirstStrand cDNA synthesis kit (Fermentas, Lithuania) according to the manufacturer protocol. The cDNA-equivalent of 5 ng RNA was used for amplification in 384-well microtitre plates in a QuantStudio 5 Real-Time PCR System (Applied Biosystems, USA) in a final reaction volume of 10 μl containing 5μL PowerUP SYBR Green master mix (Applied BiosySDs) and 0.5 μl primers mix (Integrated DNA Technologies, Singapore). All cDNA samples were amplified in triplicate. Cycle threshold (Ct) values for individual reactions were determined using QuantStudio 5, v1.4.1 data processing software (Applied Biosystems). For data analysis, Ct-value of gene of interest was first normalized against geometric mean of housekeeping genes Ct-value by the following equation [ΔCt(Gene) = Ct(Gene) – Ct(geometric mean of Tbp and Ywhaz)]. Relative mRNA level was calculated as 2-ΔCt(Gene). Relative fold change for mRNA level was calculated as 2-ΔΔCt(Gene), in which ΔΔCt(Gene) = ΔCt(Gene) of samples from different experimental condition – average of ΔCt(Gene) of controls]. Primer sequences for detection of mouse and human genes of interest are presented in Supplementary Table S1 and S2, respectively.

### Immunoblotting

Cells were washed twice with cold PBS and lysed within the wells with RIPA buffer (Thermo Scientific, 89900), supplemented with 1% protease inhibitor cocktail (Sigma-Aldrich, P8340) and 1% phosphatase inhibitor cocktail (Sigma-Aldrich, P0044). Cell lysates were sonicated prior to protein quantification. Total protein concentration was measured using Pierce 660 nm Protein Assay Reagent (Thermo Scientific, 22660) according to manufacturer’s protocol. Equal amounts of protein were loaded onto 8% or 10% denaturing SDS-polyacrylamide gels. Electrophoresis was performed to resolve proteins based on their molecular weight. Resolved proteins were electrotransferred onto polyvinylidene difluoride (PVDF) membrane (GE Healthcare, USA) at 100V. Membranes were blocked with 5% BSA in 1x Tris-buffered saline (TBS) for one hour at room temperature and incubated with antibody against mouse CEBPα, HSP90, PPARγ, pSMAD2, SMAD2, pSMAD3 and SMAD3 (Supplementary Table S3) overnight at 4°C in 5% BSA, 1x TBS, supplemented with 0.1% Tween-20. Membranes were washed three times with 1x TBS and incubated with horseradish peroxidase conjugated, goat anti-rabbit IgG secondary antibodies (1:5000, Thermo Scientific, 31460) in 5% BSA, 1x TBS, supplemented with 0.1% Tween-20. Membranes were washed three times with 1x TBS. Signals on membrane were visualized with enhanced chemiluminescence (ECL) substrate (Cyanagen, Italy, XLS142,0250) and iBright FL1500 Imaging System (Invitrogen). The intensity of bands was quantified using iBright Analysis Software V4.0.1 (Invitrogen).

### Statistical analysis

All statistical analyses were performed with GraphPad Prism (Version 8; GraphPad Software, Inc., USA). Differences were considered significant when probabilities (P) were less than 0.05.

## Acknowledgements

We thank Annalena Moliner, Eunice Sim and Ket Yin Goh for technical advice and assistance. This research was funded by grants to C. F. I. from the Singapore National Medical Research Council (NMRC/CBRG/0107/2016), ODPRT of the National University of Singapore (Aspiration Fund Partner), and Swedish Research Council (2016-01538 and 2020-01923).

## Author contributions

E.-S. L. performed all experiments except studies on *Ap2*^CRE^::*Acvr1b*^fx/fx^ mutant mice (performed by T. G.) and *AdipoQ*^CRE-ERT2^::*Acvr1b*^fx/fx^ mutant mice (performed by R. K. S.). A.S. provided human adipose tissue samples. E.-S. L. and C. F. I. designed and planned the experiments. E.-S. L. wrote the first draft of the manuscript and figures. C. F. I. wrote the final version of the manuscript and figures.

## Data availability

All data are contained within the manuscript.

## Supplementary Information

**Supplementary Table S1.**
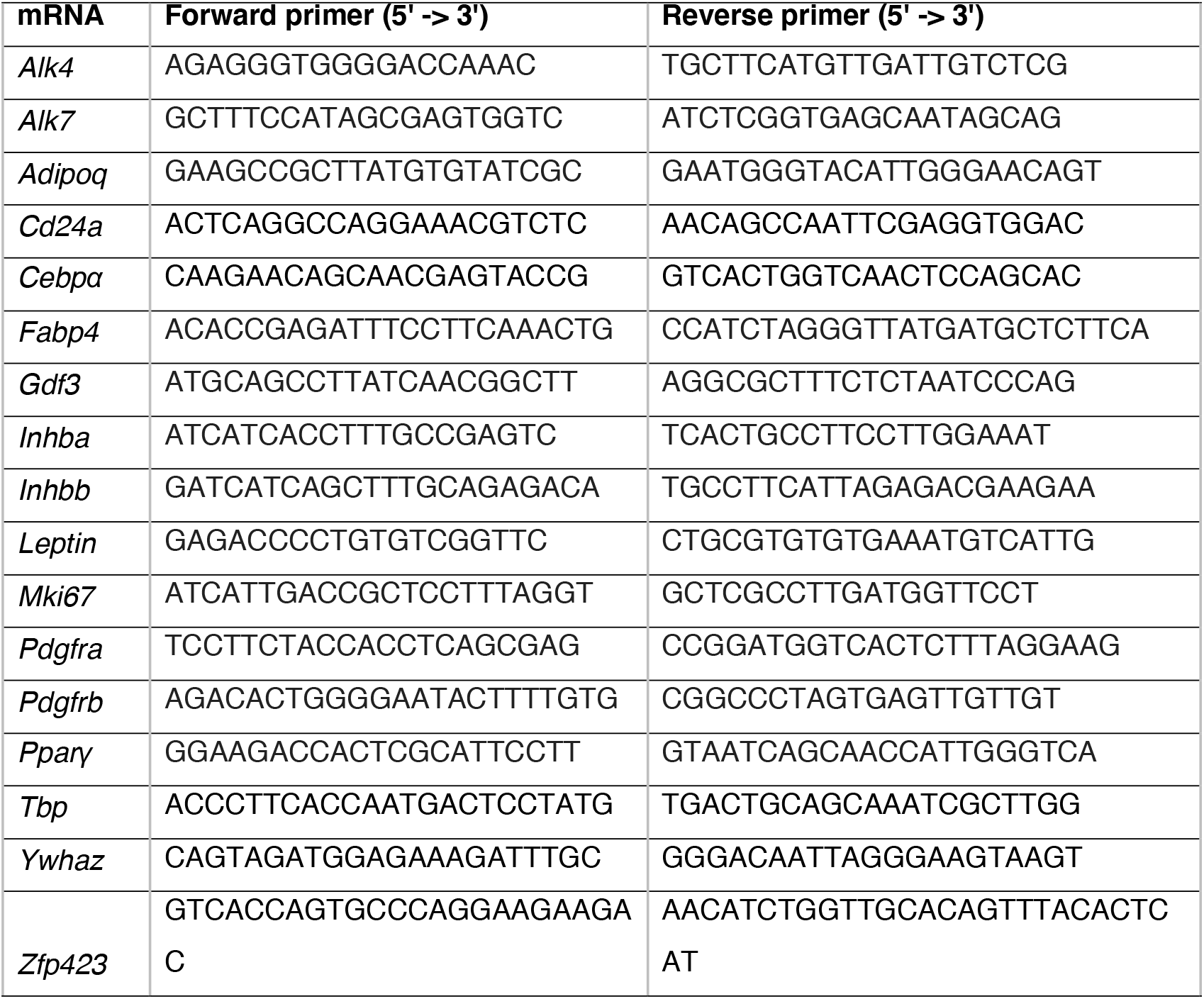
Primer sequences of mouse genes.

**Supplementary Table S2.**
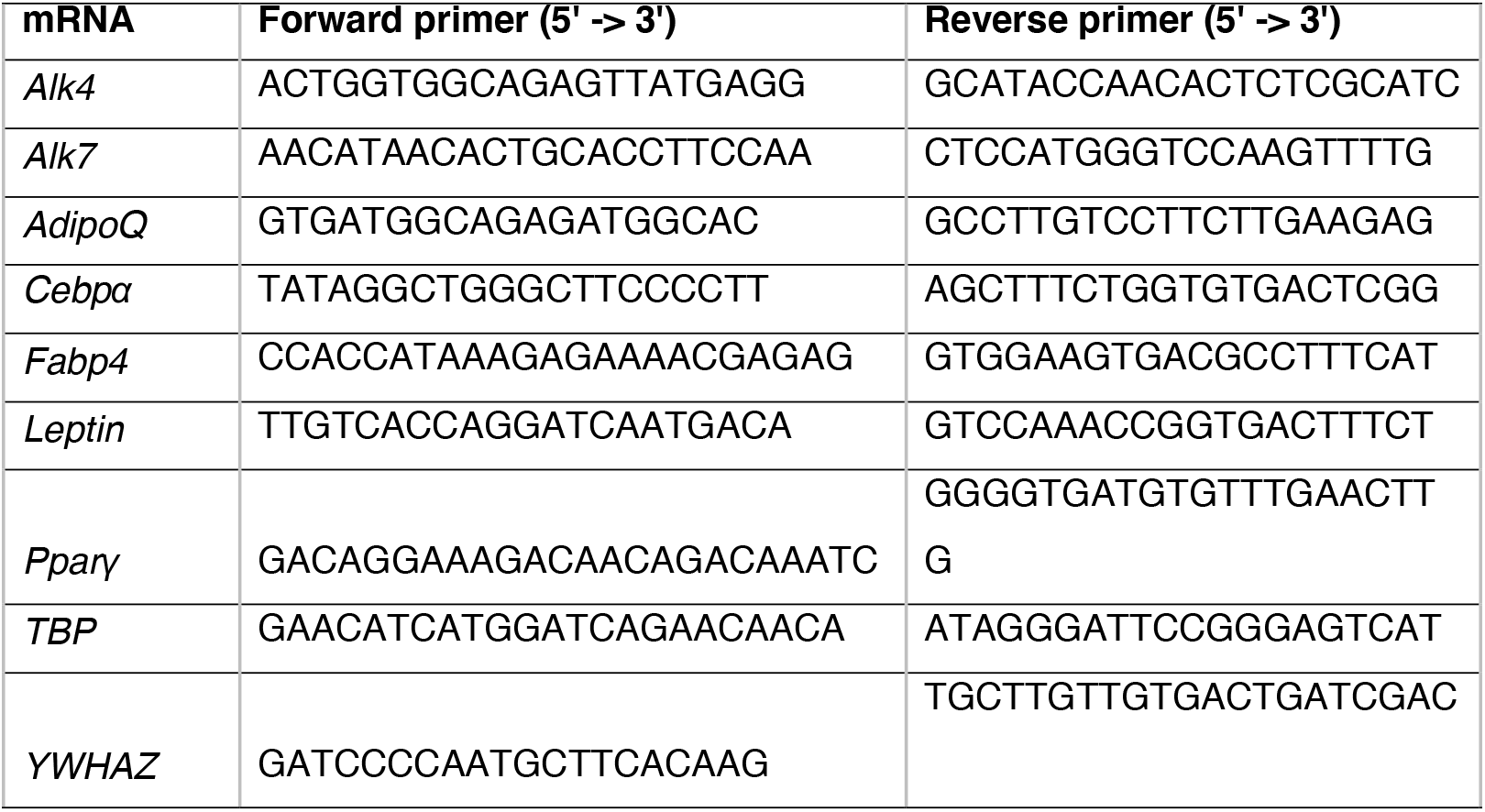
Primer sequences of human genes.

**Supplementary Table S3.**
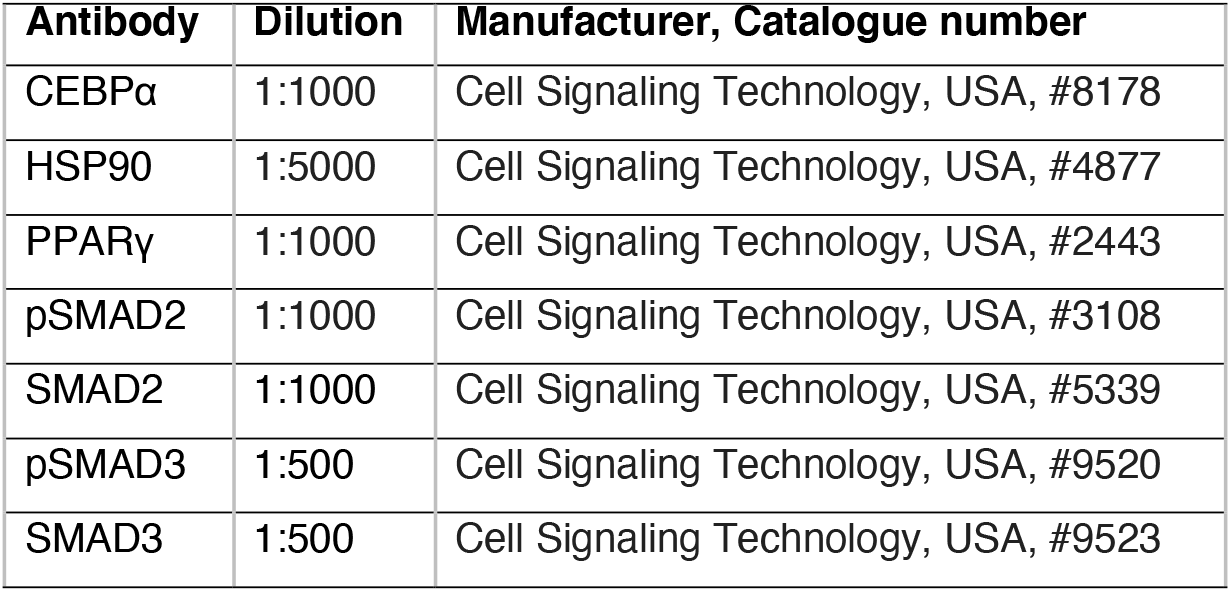
Antibodies used for immunoblotting.

| **Antibody** | **Dilution** | **Manufacturer, Catalogue number** |
| --- | --- | --- |
| CEBPα | 1:1000 | Cell Signaling Technology, USA, #8178 |
| HSP90 | 1:5000 | Cell Signaling Technology, USA, #4877 |
| PPARγ | 1:1000 | Cell Signaling Technology, USA, #2443 |
| pSMAD2 | 1:1000 | Cell Signaling Technology, USA, #3108 |
| SMAD2 | 1:1000 | Cell Signaling Technology, USA, #5339 |
| pSMAD3 | 1:500 | Cell Signaling Technology, USA, #9520 |
| SMAD3 | 1:500 | Cell Signaling Technology, USA, #9523 |

